# Evidence for prereg posters as a platform for preregistration

**DOI:** 10.1101/833640

**Authors:** Kimberly Brouwers, Anne Cooke, Christopher D. Chambers, Richard Henson, Roni Tibon

**Affiliations:** School of Medicine, Medical Sciences and Nutrition, University of Aberdeen, UK, AB24 3FX; British Neuroscience Association, The Dorothy Hodgkin Building, University of Bristol, Whitson Street, Bristol, UK, BS1 3NY; Cardiff University Brain Research Imaging Centre (CUBRIC), School of Psychology, Cardiff University, UK, CF10 3AT; MRC Cognition & Brain Sciences Unit, University of Cambridge, UK, CB2 7EF; Department of Psychiatry, University of Cambridge, UK, CB2 0SZ

## Abstract

Prereg posters are conference posters that present planned scientific projects. We provide preliminary evidence for their value in receiving constructive feedback, promoting open science, and supporting early career researchers.

---

In recent years, it has been repeatedly shown that the results of many scientific studies fail to replicate^1,2,3^. The growing awareness of this problem has prompted several attempts to tackle it. One way is to submit a *registered report* to a journal, which is peer-reviewed and has the opportunity to be revised before any data are collected^3,4,5,6^. Not only does the peer-review potentially improve the study design and analyses, but once accepted, the design and analyses are “locked”, preventing researchers from adjusting their hypotheses or analyses post hoc, i.e., after having seen the data (reporting such additional analyses is allowed, but only when transparently labelled as “exploratory”). Moreover, the journal is committed to publishing the results, whatever the outcome, which reduces the “file-drawer” problem of many null results not being published^7^. Registered reports, first offered by the journal *Cortex*, are now supported by over 200 peer-reviewed journals, including *Nature Human Behaviour*.

Another approach is to *preregister* studies on a publically-accessible platform such as the Open Science Framework^6,8^. Though they can be locked, such preregistered studies are not normally peer-reviewed prior to data collection, and they are not binding for any subsequent publications. Indeed, substantial deviations from preregistered plans have been observed^9^. Two likely reasons for the large number of deviations are immature preregistration (e.g., of a protocol that was not yet finalised) and acceptance of post-preregistration feedback (e.g., following presentation of data at a conference).

Recently, we proposed another possible avenue to help preregistration, namely presenting posters at academic conferences about planned research, before data are collected^10^; so-called “prereg posters”. This allows presenters to receive feedback on their hypotheses, design and analyses from their colleagues (conference attendees), which is likely to improve the study. In turn, this can improve more formal preregistration, reducing the chances of subsequent deviation, and/or facilitate submission of the work as a registered report. Moreover, colleagues with shared scientific interests become aware of the study early on, which can open the door to collaborations.

Prereg posters were recently adopted by the BNA*2019* Festival of Neuroscience (https://meetings.bna.org.uk/bna2019/), a biennial event organised by the British Neuroscience Association (BNA). Such posters have also been allowed by at least three other conferences since then: FLUX (https://fluxsociety.org/2019-new-york), BACN (https://www.bacn.co.uk/conferences) and ICON (https://www.helsinki.fi/en/conferences/international-conference-of-cognitive-neuroscience-2020). The BNA*2019* Festival organisers additionally collected informal survey data about prereg posters, which we report here as preliminary evidence for their value.

Nearly a fifth (100/491) of all submitted posters at the BNA*2019* Festival conformed to the new prereg format, covering a diverse range of neuroscience topics and disciplines, and the overall impression was that they were enthusiastically welcomed. The informal survey data were collected at two time points (before and after the event), via two different online surveys. The full set of questions and response data are available here: https://osf.io/3h6w8/. The first informal survey (pre-conference survey) was administered three months prior to the event, and offered to all 445 participants whose poster was accepted. Of the 200 who responded, 151 were presenters of traditional posters and 49 of prereg posters. The second informal survey (post-conference survey) was administered one month after the event, and was completed by 95 participants, 66 of whom were presenters of traditional posters, and 29 of prereg posters. Below, we present data from the questionnaire items that address three main themes: prereg posters as means to (1) receive valuable feedback, (2) promote open science, and (3) support early career researchers (ECRs).

## Getting valuable feedback

As a presenter, the motivation to present a prereg poster is clear: it allows one to get feedback on work at early stages. Indeed, 33% of the 123 responses to the pre-conference question “why did you submit a prereg abstract instead of a traditional abstract?” indicated that this choice was made in order to get feedback (either in general or on specific analyses) before completing the research. As shown in Figure 1, the groups differed in the type of feedback that they subsequently received at the conference. Interestingly, presenters of traditional posters mostly received feedback regarding future projects, whereas presenters of prereg posters mostly received feedback regarding methods and experimental design.

**Figure 1:**
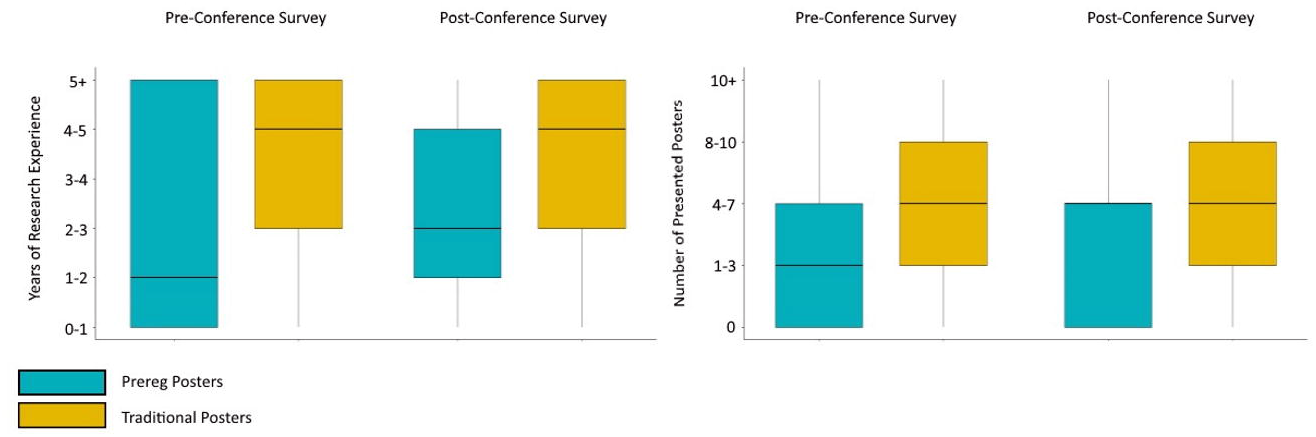
Kind of feedback received for posters. Distribution of responses to the post-conference survey question “what kind of feedback did you receive”, among presenters of prereg posters (left) and traditional posters (right). Nr represents the total number of responses.

Furthermore, the groups gave similar estimations of the likelihood of the presented research being published in a peer-reviewed journal upon completion (prereg: *Median* = 6 [scale: 1 – “highly unlikely” to 7 – “highly likely”], *IQR* = 2; traditional: *Median* = 6, *IQR* = 2). This is even though projects presented as prereg posters were at their early stages, and in most cases were still prior to data collection. While it could be the case that researchers’ estimation of publication likelihood remains stable throughout the course of a project, we speculate that it could also reflect the presenters’ trust in the preregistration system, where publication does not depend on results, and/or the presenters’ impression that they received valuable feedback that would promote their ability to publish the work later.

One potential objection to prereg posters is that conference attendees may not bother to discuss “half-baked” work and therefore there is no point in “wasting” conference space on posters that would not attract visitors. However, responses to two questions in the post-conference survey—“did you receive feedback on your work during the poster session?” and “how many people talked to you about your poster during the poster session?”—suggest that this is not the case: prereg posters did not receive less feedback than traditional posters (prereg = 75%; traditional = 68%), nor fewer visitors (prereg: *Median category* = 2 [5-8 people], *IQR* = 1; traditional: *Median Category* = 2 [5-8 people], *IQR* = 1), suggesting that conference attendees do not avoid prereg posters, despite the fact that these posters present planned or preliminary work.

## Promoting open science

Our second theme concerns attitudes among the participants towards open science in general, and preregistration in particular. Two questions from the post-conference survey were indicative of such attitudes: (1) “to what degree would you say that preregistration of work is necessary in today’s neuroscience community?” and (2) “to what degree did BNA*2019* increase your awareness of preregistration?”. In general, presenters of both prereg and traditional posters agreed that preregistration is necessary (prereg: *Median* = 4 [scale: 1 - “not at all” to 5 – “incredibly necessary”], *IQR* = 1; traditional: *Median* = 4, *IQR* = 2). Moreover, participants in both groups indicated that the event increased their awareness of preregistration, and the increment was somewhat greater for prereg poster presenters (prereg: *Median* = 6 [scale: 1 - “not at all” to 7 – “completely”], *IQR* = 2; traditional: *Median* = 5, *IQR* = 2.75). Furthermore, in the pre-conference survey, 61% of prereg posters submitters indicated that one reason they chose this form of presentation was to gain experience and confidence with preregistration and with Registered Reports. These results suggest that the attendees of BNA*2019* acknowledged the importance of preregistration and that the event was helpful in promoting this understanding further, particularly for those presenting prereg posters.

## Supporting early career researchers

Although all scientists can benefit from their colleagues’ comments, these can be particularly beneficial to ECRs who are still developing their networks and expertise in the field. Because on most occasions (especially for ECRs), travel funding is only available when work is presented, and given that ECRs often have fewer existing datasets to present compared to more senior colleagues, prereg posters can provide ECRs additional opportunities to attend academic events. Therefore, the opportunity to present a prereg poster may be especially compelling for ECRs. Data from the present informal surveys that are relevant to this question are shown in Figure 2. While the data do not support strong claims regarding the relations between career stage and the choice to present a prereg poster, they do indicate that presenters of BNA2019 prereg posters had more limited research experience and had presented fewer posters throughout their career compared to traditional poster presenters.

**Figure 2.**
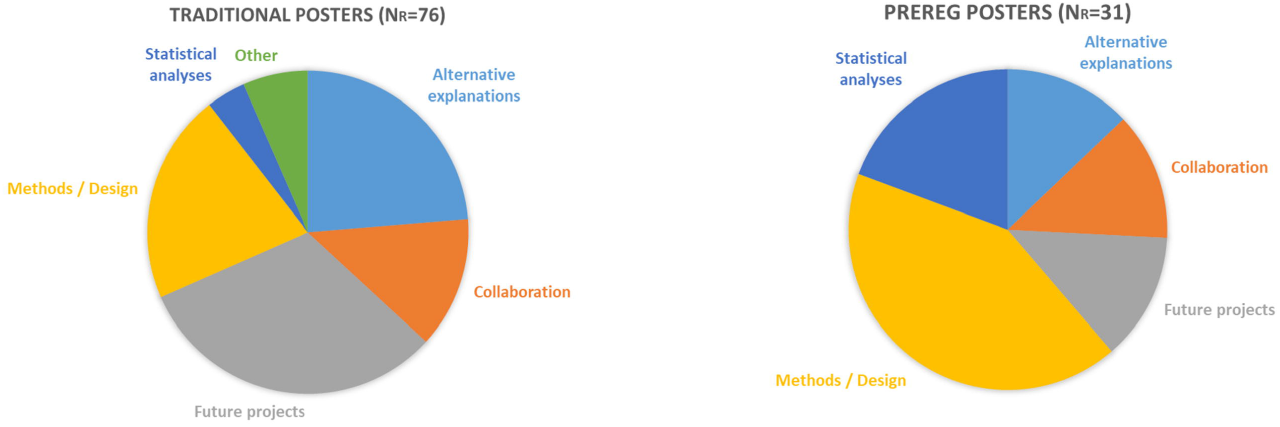
Indications of presenters’ career stage. Distribution (around the median) of the years of research experience (left) and number of presented posters in academic conferences (right) among the presenters of prereg posters (cyan) and traditional posters (amber). Within each plot, boxplots on the left represent data obtained from the pre-conference survey (N=200) and boxplots on the right represent data obtained from the post-conference survey (N=95)

## Conclusions and outlook

The qualitative evidence reviewed above supports our prior claim that prereg posters can be a useful tool in promoting academic discussion of planned and on-going research, encouraging open science, and benefiting early career researchers. We hope this encourages the adoption of prereg posters by an increasing number of future scientific conferences. Notably, unlike some forms of study registration, whose main goal is to commit researchers to a registered plan, the main goal of prereg posters is to provide an opportunity for adjusting and improving studies before they are formally registered, following feedback from colleagues. Prereg posters can be routinely archived in a public repository, and then cited in any subsequent formal preregistrations that arise from the nascent protocol. This would capture the history and thus provenance of the idea generation. Indeed, it may be helpful to distinguish between “unlocked preregistration”, which refers to research plans that are presented as posters or uploaded to a public website, but are still subject to change, and “locked (pre)registration”, such as a registered report, which refers to a finalised research plan.

As we previously discussed^10^, and as with any new initiative, we believe it is vital to instruct all the people involved in conferences (organisers, presenters, poster reviewers and attendees) about the aim and value of these posters, otherwise the initiative might flounder (e.g, if reviewers score prereg posters lower owing to their lack of results, because the reviewers were not properly informed of the aims). Additionally, at least until they become more customary, we recommend highlighting prereg posters in the conference program and at the stand (e.g., by open science badges, as done at BNA*2019*). Further practical advice and useful tips can be found on the BNA website: www.bna.org.uk/mediacentre/news/pre-reg-posters/. Importantly, we believe that conferences and academic events should not only support specific scientific topics, but also act as venues that increase the quality of scientific research.

## Acknowledgements

Authors KB and AC are/were employed by the British Neuroscience Association (BNA) and its credibility in neuroscience programme, supported by the Gatsby Charitable Foundation (GAT3674), and played a key role in the design, conceptualization, data collection, analysis, and preparation of this manuscript. The BNA further provided the venue for data collection (at the BNA2019 Neuroscience Festival). Other funders contributed to the time of the authors, namely a British Academy Postdoctoral Fellowship (SUAI/028 RG94188) to RT, UK Medical Research Council grant (SUAG/010 RG91365) to RH, and ERC Consolidator grant (647893-CCT) to CDC, but had no role in the conceptualization, data collection and analysis, decision to publish or preparation of the manuscript.

## Author contribution

K.B: designed, planned, and carried out the survey; collected the data; provided input on the paper. A.C: conceived the idea; designed and planned the survey; provided input on the paper. C.D.C: provided input on the paper. R.H: directed the project together with R.T; commented on structure and analysis approach since early drafts; provided input on the paper. R.T: directed the project together with R.H; analysed the data; wrote the paper with input from all authors; shared the data on OSF.

